# Stereotypic persistent B cell receptor clonotypes in Alzheimer’s Disease

**DOI:** 10.1101/2023.09.07.554570

**Authors:** Hyunji Yang, Namphil Kim, Yonghee Lee, Duck Kyun Yoo, Jinny Choi, Ki Woong Kim, Jong Bin Bae, Ji Won Han, Sunghoon Kwon, Junho Chung

**Author notes:** **Corresponding author** S. Kwon or J. Chung.

## Abstract

We constructed B cell receptor (BCR) repertoires *in silico* using peripheral blood (PB) samples collected from 44 Alzheimer’s Disease (AD) patients at baseline and 37 patients at follow-up. For the control group (CG), we used BCR repertoire data from the chronologically collected PB samples of 55 healthy volunteers vaccinated with SARS-CoV-2 mRNA. The AD patients shared 3,983 stereotypic non-naïve BCR clonotypes not found in CG, and their degree of overlap between patient pairs were significantly higher than that of CG pairs, even with the SARS-CoV-2 spike protein triggering a concerted BCR response. Twenty stereotypic non-naïve AD patient-specific BCR clonotypes co-existed in more than four patients and persisted throughout two sampling points. One of these BCR clonotypes encoded an antibody reactive to the Aβ42 peptide. Our findings strongly suggest that AD patients are exposed to common (auto)antigens associated with disease pathology, and their BCR repertoires show unique signatures with diagnostic potential.

## Introduction

Alzheimer’s disease (AD) is a progressive neurodegenerative disorder that causes dementia, cognitive decline, memory loss, and impaired daily functioning. Thus far, the presence of amyloid beta peptide (Aβ) plaques and neurofibrillary tangles (NFTs) composed of phosphorylated tau proteins within the central nervous system (CNS) have been recognized as major contributors to neuroinflammation and the pathological hallmarks of AD^1, 2^. However, recent studies have revealed that the pathogenesis of AD is not solely restricted to these changes in the CNS^3^. An increasing body of evidence points to the peripheral immune system as another major player intimately linked to AD pathology^4-7^. Several studies have demonstrated that the peripheral homeostasis of T lymphocytes is disrupted in AD patients^8-10^, but little is known about the role of peripheral B lymphocytes in AD. A single-cell RNA sequencing analysis found that the progression of AD is associated with a reduction in the number of B cells in the peripheral blood, as well as differentially expressed genes in B cells^11^. B cells play a fundamental role in the adaptive immune system particularly in humoral immunity, by recognizing antigens via membrane-expressed B cell receptors (BCR) and producing antigen-specific immunoglobulin (Ig) in response^12^. Accordingly, recent studies have investigated the differences in immune cell type compositions and the clonal diversity of BCR repertoires among AD patients through single-cell RNA sequencing data or single-cell BCR libraries of peripheral blood mononuclear cells (PBMCs)^5,13^. The results showed that the proportion of B cells and the clonal diversity of both the heavy and kappa (κ) light chains of BCR repertoires all significantly decreased with the progression of AD. Previous reports including ours have characterized the BCR repertoires of AD patients, such as isotype distribution, variable (V) and joining (J) gene usage, somatic hypermutation (SHM) patterns, and diversity of the complementarity-determining region 3 (CDR3)^5,14^. Despite the success in highlighting the importance of BCRs in AD neuropathology, the findings were limited by the relatively small sample sizes of 3 and 10 and also lack of chronological sampling. Furthermore, they have not characterized the stereotypic BCRs at an individual sequence level or identified their specific antigens.

In this study, we identified BCR clonotypes that are persistently present and shared among AD patients, but not in healthy populations. For this analysis, we collected the peripheral blood (PB) of 44 AD patients at baseline and chronologically from 37 patients at follow-up with an interval of 14 - 53 weeks, and re-constituted their BCRs *in silico*. For the control group (CG), we used BCR repertoires from 55 healthy volunteers injected with the severe acute respiratory syndrome coronavirus 2 (SARS-CoV-2) mRNA vaccine at baseline. Their PB samples were collected a maximum of six times over 42 weeks (285 PB samples obtained chronologically)^15^ and were expected to share a high number of SARS-CoV-2 vaccine-induced BCR clonotypes.

In addition, one of the identified stereotypic AD patient-specific non-naïve BCR clonotypes was found to encode antibodies reactive to the Aβ42 peptide, a well-known key antigen associated with AD progression^16^.

## Main text

### Selection of stereotypic AD patient-specific non-naïve BCR clonotypes

For a period of 75 weeks, PB samples were collected from 44 AD patients with an average age of 75 years (standard deviation; 7.04) and sex ratio of 34% and 66% (male vs female). PB samples were collected twice with an interval of 14 to 53 weeks, with the exception of seven patients who withdrew from the study after sampling at time point (TP) 1 (Patient ID 3, 5, 8, 19, 28, 30, and 37) (Fig. 1a). The duration of the first PB sampling period was 61 weeks, and the second PB sampling 34 weeks, respectively. From these 81 PB samples, the cDNA was prepared and the gene fragments encoding both the variable domain (V^H^) and the N-terminal region of the BCR heavy chain constant domain 1 (CH1) were subjected to amplification using a specific primer set (Extended Table 2). The amplified products were then analyzed by next-generation sequencing (NGS) with a median read of 245,033 (standard deviation of 184,462, Extended Table 3). For BCR repertoire analysis, we first removed naïve BCR sequences, which we define as those with the IgM or IgD isotypes containing less than or equal to 1 SHM (Step 1, Fig. 1c). We then grouped the remaining sequences into clonotypes, which we defined as a collection of sequences encoded by identical IGHV and IGHJ genes and the same CDR3 sequence at the amino acid level^17^. Among these non-naïve BCR clonotypes, AD patient-specific clonotypes were selected by excluding the clonotypes also found in our prior studies among non-AD individuals (Step 2, Fig. 1c). This non-AD CG was constituted with a total of 55 healthy subjects injected three times with the SARS-CoV-2 vaccine from our previous study^15^. From this group, 285 PB samples were obtained chronologically with a minimum of 35,000 functional BCR sequences^15^ (Fig. 1b, Extended Table 3). Then the stereotypic clonotypes shared by more than two AD patients were selected (Step 3, Fig. 1c). Before further analysis, we excluded stereotypic clonotypes that failed to show different BCR sequences at the nucleotide level in two or more AD patients, as they were likely to be false positives by aerosol molecular contamination and index switching during NGS library preparation and sequencing (Step 4, Fig. 1c)^15^. The described procedures were then replicated in the CG group to collect the control stereotypic non-naïve BCR clonotypes. At the end of this selection process, we were able to identify 3,983 stereotypic non-naïve BCR clonotypes specific to the AD patient group, and 83,821 specific to the CG, respectively (Fig. 1c).

**Figure 1.**
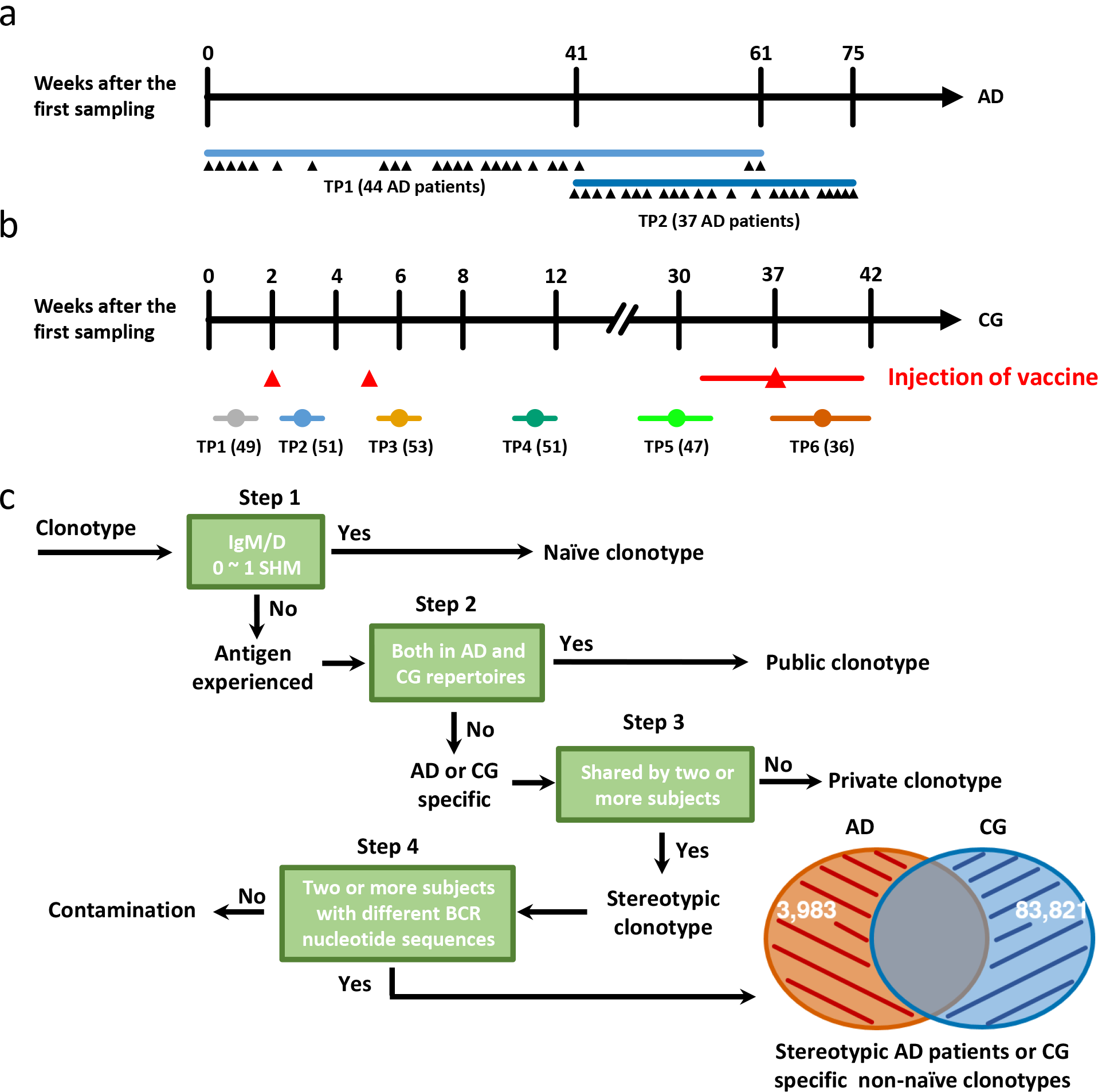
Peripheral blood sampling and BCR repertoire analysis. a, From AD patients, PB samples were collected twice with an interval of 14 to 53 weeks, with the exception of seven patients who withdrew from the study after time point (TP) 1 sampling (Patient ID 3, 5, 8, 19, 28, 30, and 37). The black arrowheads represent the exact sampling points in the given time frame. b, The control group (CG) comprises of 55 vaccine recipients who were subjected to peripheral blood sampling before their first dose of vaccination (TP1, 49 subjects), 1 week after the 1^st^ dose (TP2, 51), 1 week after the 2^nd^ dose (TP3, 53), 6 weeks after the 2^nd^ dose (TP4, 51), 30 weeks after the 2^nd^ dose (TP5, 47), and 1 to 4 weeks after the 3^rd^ dose (TP6, 36). c, BCR clonotypes were assembled from the BCR repertoires of the AD patients and CG, with BCR clonotypes being defined as a group of BCR sequences encoding identical IGVH and IGVJ genes and the same CDR3 sequence at the amino acid level. If a clonotype contained only naïve sequences, which were defined as those with IgD or IgM isotypes and 0 – 1 somatic hypermutations (SHM), the clonotypes were excluded from further analysis. The clonotypes present in both the AD patient group and CG were excluded, and the stereotypic clonotypes present in more than one AD patient or more than one CG subject were selected. To exclude the possibility of molecular contamination during NGS library preparation or NGS, we confirmed at least two AD patients or CG subjects with different BCR sequences at the nucleotide level in each identified clonotype. We discarded all clonotypes that failed to meet this criteria. Finally, we obtained 3,983 AD patient-specific and 83,821 CG subject-specific non-naïve BCR clonotypes.

### Stereotypic AD patient-specific non-naïve clonotypes showed a significantly higher degree of overlap in patient pairs compared to CG pairs

We then analyzed the distribution of stereotypic AD patient-specific non-naïve BCR clonotypes in 1,892 ((44 X 44)-44) and 1,332 ((37 X 37)-37) patient pairs in the first and second TPs of PB sampling, respectively. We calculated the degree of overlap for stereotypic non-naïve BCR clonotypes in each pair using the cosine similarity method, with one-hot encoded vectors representing the presence or absence of each stereotypic non-naïve clonotype in a patient’s BCR repertoire. We also obtained the degree of overlap for the CG pairs over the six PB sampling TPs. As expected, there was a dramatic increase in the average cosine similarity compared to the pre-vaccinated state (TP1) at TP3, TP4, with the highest increase at TP6. Each TP corresponds to 1 week after the second-(TP3), 6 weeks after the second-(TP4), and 1 to 4 weeks after the third-injection (TP6) of the SARS-CoV-2 vaccine. This increase in average cosine similarity can be attributed to the concerted BCR responses toward the SARS-CoV-2 immunogen (Fig 2a-c, p = 4.92 × 10^−6^, p = 2.52 × 10^−30^, p = 2.26 X 10^−112^). The AD patient pairs also showed significantly higher average cosine similarity at both TPs compared to the CG’s baseline (TP1) (p = 2.06 X 10^−21^, p = 1.22X10^−169^). The cosine similarity value of the AD patient samples was higher at TP2, possibly due to the shorter PB sampling duration (61 weeks vs 34 weeks), which would provide less diverse seasonal antigens to AD patients. Unexpectedly, the average cosine similarity of the AD patients’ TP2 stereotypic non-naïve BCR clonotypes was significantly higher than even that of the CG at TP6 (p = 1.86x10^−6^). This observation is interesting especially under the consideration that the PB sampling duration was much longer in the AD patients compared to the CG (34 weeks vs 6 weeks). To account for the fact that we are not aware of the SARS-CoV-2 infection history or vaccination status of the AD patients, we performed the same analysis on all stereotypic non-naïve BCR clonotypes identified in the two groups without excluding the stereotypic non-naïve BCR clonotypes observed in the other group (excluding step 2, Fig. 1c). The resulting cosine similarity values of the AD patients’ stereotypic non-naïve BCR clonotypes at TP1 and TP2 were both significantly higher than those of the CG at all TPs except for TP6, which did not show statistically significant difference (Extended Figure 1a-c). These results clearly demonstrate that AD patients share stereotypic non-naïve BCR clonotypes at a degree compatible to CG at TP6, irrespective of the AD patients’ longer PB sampling period.

**Figure 2.**
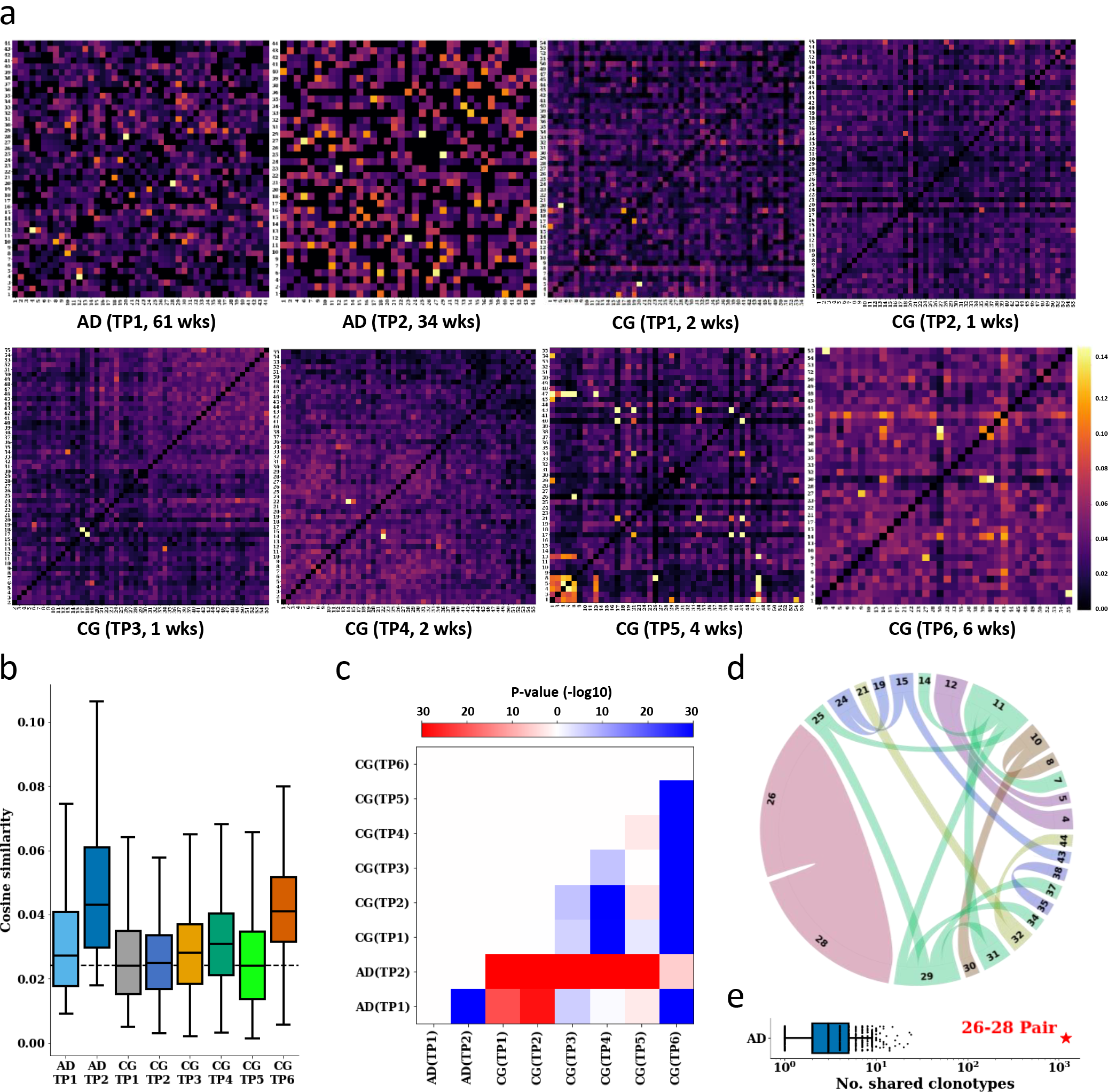
The sharing degree of stereotypic non-naïve BCR clonotypes among AD patient and CG subject pairs. a, Heat maps show the sharing degree of stereotypic AD patient- or CG subject-specific non-naïve BCR clonotypes in AD patient or CG subject pairs. The cosine similarity between two one-hot encoded linear vectors representing the presence (1) or absence (0) of each stereotypic BCR clonotype. PB sampling TP and its duration are denoted inside parentheses. b, Boxplots representing the distribution of cosine similarity for all AD patient or CG subject pairs at each PB sampling TP. The dashed line represents the median cosine similarity value of CG subject pairs at TP1. c, A heat map representing the P-values about the difference in their cosine similarity value in AD patient or CG subject pairs at every PB sampling TPs. The color of the heat maps indicate whether the median value on the y axis is larger (red) or smaller (blue) than that on the x axis. The intensity of said colors is encoded by the P-value. d, Chord plot displaying the similarity matrix of the top 20 AD patient pairs. The similarities were calculated from the combined repertoires of both time points if applicable. The chord width represents the similarity value. The thickest chord connects patient 26 and 28 and represents the value of 0.961, while the thinnest chord connects patients 11 and 25 and represents a value of 0.063. e, The number of stereotypic AD patient specific non-naïve BCR clonotype shared by each AD patient pair. The pair with the highest number of shared clonotypes is denoted with a red star.

Our analysis also showed that the degree of overlap in the stereotypic AD patient-specific non-naïve BCR clonotypes significantly differs among patient pairs. This suggests that AD patients can be placed in subgroups categorized by their BCR clonotypes. To assign these patient subgroups, the stereotypic AD patient-specific non-naïve BCR clonotypes at all TPs in each patient were combined, and the cosine similarity of patient pairs were calculated. Twenty patient pairs with the highest cosine value were selected and displayed on a Chord plot (Fig. 2d), in which the patients were divided into six subgroups. Among these pairs, one patient pair (patient ID 26 and 28) had an outlying cosine similarity value of 0.961, and the number of BCR clonotypes was also noticeably higher than the other pairs (Fig. 2e). Indeed, between the 1,225 and 1,228 stereotypic non-naïve BCR clonotypes in patient ID 26 and patient ID 28, respectively, 1,179 clonotypes were shared. The high degree of overlap in the BCR clonotypes within these AD patient subgroups strongly suggests the presence of common and possibly auto-antigens within the subgroup.

### A highly shared and persistent AD patient-specific non-naïve BCR clonotype encodes an antibody reactive to the Aβ42 peptide

To identify the antigens to which these stereotypic AD patient-specific non-naïve BCR clonotypes are reactive, we selected twenty clonotypes that were shared between four or more patients (Fig. 3a). We also analyzed the persistence of these clonotypes over the two TPs. One clonotype encoded by the IGHV3-7/IGHJ1 genes with the CDR3 sequence VRLAEYFQN was present at both TPs in three patients. When we allowed one amino acid mismatch in the CDR3 sequence, we found an additional patient (patient ID 34) with a similar clonotype (CDR3 sequence: VRLAEYFQH) (Fig. 3a). In the 285 PB samples of the CG, we did not find any BCR clonotypes encoded by the IGHV3-7/IGHJ1 genes and with a CDR3 sequence homologous to VRLAEYFQN, with one amino acid mismatch allowance (Fig. 3b). We then displayed all the BCR clonotypes encoded by IGHV3-7/IGHJ1 with CDR3 sequences homologous to VRLAEYFQN, given one amino acid mismatch allowance (Fig. 3c). Class switch recombination (CSR) processes to IgG1 and IgG3 were found in three of four total patients, and the BCR sequences harbored 4-19 SHMs in the CDR1, CDR2, and framework regions. As CSR and SHM accumulation in BCR sequences are recognized as the hallmarks of chronic repeated exposure to antigens^18^, it is possible that these clonotypes were developed by persistent exposure to (auto)antigens.

**Figure 3.**
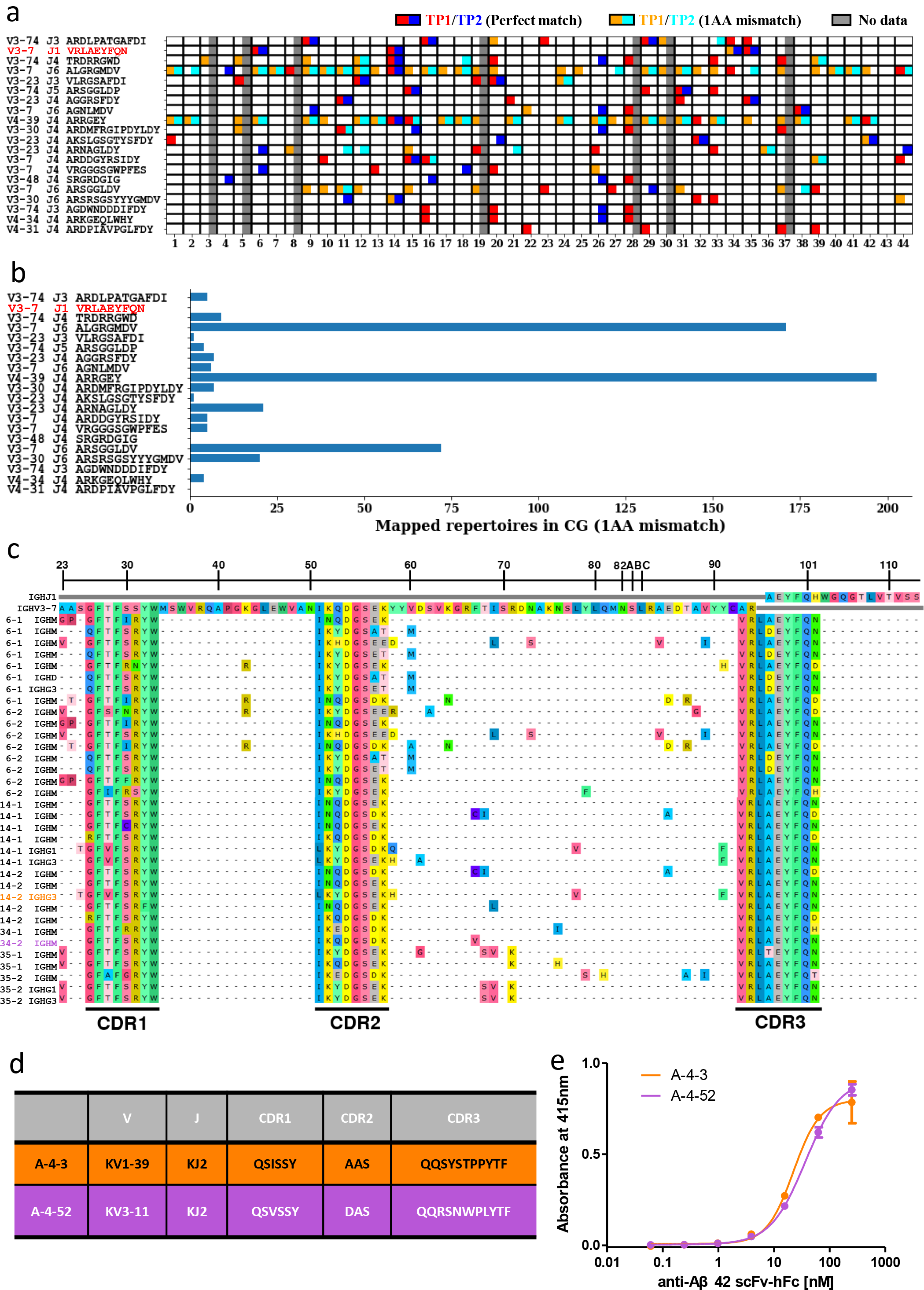
Selection of stereotypic non-naïve AD patient-specific BCR clonotypes and their characterization. a, Stereotypic non-naïve AD patient-specific BCR clonotypes shared by more than four patients were displayed. The grey squares present blank data due to the lack of PB samples. For homologous clonotypes encoded with the identical IGHV and IGHJ genes and possessing a similar CDR3 sequence given one amino acid mismatch allowance, we labeled the square similar but different colors. b, The number of CG BCR repertoire with clonotypes homologous to those displayed in a (encoded with the identical IGHV and IGHJ genes and possessing a similar CDR3 sequence given one amino acid mismatch allowance). c, BCR sequences of stereotypic non-naïve AD patient-specific BCR clonotypes clonotypes encoded by IGHV3-7/IGHJ1 genes and with VRLAEYFQN CDR3 sequence and its homologous clonotypes with one amino acid mismatch allowance at CDR3. d, The light chain sequence of the Aβ42 peptide binding scFv clones discovered from a phage display library. e, Binding of recombinant scFv-hFc fusion proteins to Aβ42 peptide coated on a microtiter plate. The absorbance difference between wells coated with Aβ42 peptide and those only blocked with BSA solution are displayed.

To further investigate this homologous clonotype, we synthesized four BCR heavy chain genes (Extended Table 4) to construct a single-chain variable fragment (scFv) phage display library, using the amplified light chain genes of AD patients (patient ID 6, 14, and 34) possessing the IGHV3-7/IGHJ1 VRLAEYFQN clonotype. We then performed bio-panning of this library on the Aβ42 peptide, a well-known autoantigen in AD^16^, and were able to find Aβ42 peptide-reactive scFv clones. When we expressed these clones as recombinant scFv-human Fc fusion proteins and performed an ELISA, two scFv clones with different kappa (κ) light chains (Fig. 3c, Extended figure 2) showed reactivity against the Aβ42 peptide, with half-maximal effective concentrations (EC50) of 73.62 nM and 86.42 nM, respectively (Fig. 3d).

Our results support that the antibodies encoded by stereotypic AD patient-specific BCR clonotypes are developed by chronic exposure to autoantigens closely related to the pathological processes of AD.

## Discussion

In this study, we found that AD patients share stereotypic non-naïve BCR clonotypes not found in CG, and the degree of their overlap between patient pairs were higher than that of CG pairs, despite the CG being exposed to the SARS-CoV-2 spike antigen to develop a very high antibody titer and concerted BCR repertoire^15^. This observation strongly suggests the presence of a common antigen among AD patients that coordinates the BCR repertoire into selected directions. When we analyzed twenty clonotypes that were shared by more than four AD patients, many of these stereotypic AD patient-specific BCR clonotypes persisted over a period of 14 to 53 weeks (Fig. 3a). In addition, when we examined IGHV3-7/IGHJ1 VRLAEYFQN clonotypes at the individual sequence level, we found not only CSR occurrences, but also the accumulation of diverse SHM profiles. We also checked the CSR status and the level of SHM in the remaining top 19 stereotypic AD patient-specific non-naïve BCR clonotypes, of which 13 showed evidence of prior CSR events, and 16 showed evidence of prior SHMs of over 10 nucleotides (Extended Table 5). These findings strongly claim that the exposure to a common antigen was likely not a single event in most cases, but rather chronic. Collectively, our results suggest the presence of a common (auto)antigen that persists in AD patients and provokes a concerted and constitutive humoral immune response.

One of the drawbacks in our study is that the age of the CG was not matched with that of AD patients. BCR repertoires are known to be influenced by the aging process^19^, and we have previously reported that many centenarians possess autoantibodies to the DNA-directed RNA polymerase II subunit RPB1, while being relatively rare in younger populations^20^. Therefore, without an age and sex matched CG, it is difficult to rule out the possibility that the evolution of these stereotypic AD patient-specific non-naïve BCRs are from the aging process rather than AD progression. In this regard, we tried to co-relate these stereotypic AD patient-specific BCR clonotypes with the pathogenic process of AD. Several autoimmune diseases were reported to show unique signatures on their BCR repertoires^21^. For example, autoantibodies to the acetylcholine receptor and muscle-specific tyrosine kinase are found among most myasthenia gravis patients and very closely associated with pathogenic processes^22^. In AD, over 70 autoantigens have been reported, to which autoantibodies were found in the blood or cerebrospinal fluid^3^. Among these autoantigens, Aβ has been recognized to play a pivotal role in AD pathogenesis. It causes synaptic impairment and neurodegeneration, thus contributing to the cognitive dysfunction observed in AD^23^. Two anti-Aβ antibodies, lecanemab and aducanumab, have been approved for clinical use in AD patients and few others are on clinical trials. For active immunotherapy, anti-Aβ vaccines capable of eliciting a polyclonal anti-Aβ antibody response are also on clinical trials (NCT05462106 & NCT03461276). In this study, we focused on the IGHV3-7/IGHJ1 VRLAEYFQN clonotype and revealed that it encodes an antibody reactive to the Aβ42 peptide, a well-known autoantigen in AD^16^. With this finding, we believe that more stereotypic AD patient-specific BCR clonotypes may be related to the pathophysiology of AD.

Over the past decade, research findings highlighting the relationship between autoantibodies and the progression of brain diseases have provoked a paradigm shift in neurological diseases, opening opportunities for novel diagnostic modalities^24^. In this study, we identified 3,983 stereotypic AD patient-specific non-naïve BCR clonotypes. It is also noteworthy that these clonotypes have been filtered by the strictest sequence contamination rationale reported so far, leaving little room for false positives. Blood-based biomarkers, like phosphorylated tau in plasma, hold great promise to revolutionize the diagnostic and prognostic work-up of AD in clinical practice^25^. BCR clonotypes are potential blood-based biomarkers and we have already reported the existence of disease-specific stereotypic BCR clonotypes among Sjogren syndrome patients^26^ as well as COVID-19 patients^18^. Additionally, the accumulating single B cell sequencing data of AD patients can be combined with our clonotype database to match the natural heavy and light chain pairs of BCR clonotypes to identify individual autoantigens, which could further facilitate the evaluation of BCR clonotypes as a blood-based biomarker. With these insights, we could also classify AD patients into subgroups based on the shared degree and types of stereotypic BCR clonotypes. A recent study reported that AD can be classified into three molecular subgroups by transcriptomics data, and the clinical features were found to differ in each group, suggesting the potential for personalized diagnosis and treatment^27^.

In summary, we found that AD patients share many BCR non-naïve clonotypes, of which homologous clonotypes are not or rarely found in healthy non-AD populations. By analyzing these commonly found stereotypic AD-patients specific clonotypes, we demonstrated that they were formed by repeated exposure to the antigens shared among the AD patients, such as Aβ. Additionally, as AD patients can be sub-grouped depending on the BCR signature, we believe that BCR repertoires have potential to become a blood-based biomarker for the improved diagnostic and prognostic evaluation of AD.

## Methods

### Study subjects

We recruited participants who had been diagnosed with mild cognitive impairment (MCI) or dementia due to Alzheimer’s disease (AD) from the dementia clinics of Seoul National University Bundang Hospital (SNUBH) and the Korean Longitudinal Study on Cognitive Aging and Dementia (KLOSCAD)^28^. We collected demographic information, Clinical Dementia Rating (CDR)^29^, and neuropsychological test scores of the Consortium to Establish a Registry for Alzheimer’s Disease Assessment Packet (CERAD-K) Neuropsychological Assessment Battery (CERAD-K-N)^30,31^ from all subjects. We also conducted brain magnetic resonance imaging (MRI) and 18F-Florbetaben positron emission tomography (PET) scans on all participants. All recruited participants were amyloid positive on their PET scans.

A panel of research neuropsychiatrists made the dementia diagnoses according to the criteria in the Diagnostic and Statistical Manual of Mental Disorders, Fourth Edition (DSM-IV)^32^. They diagnosed the Alzheimer’s disease (AD) dementia according to the criteria of the National Institute of Neurological and Communicative Disorders and Stroke and the Alzheimer’s Disease and Related Disorders Association^33^. MCI was diagnosed according to the Consensus Criteria from the International Working Group on MCI^34^. The presence of objective cognitive impairment was ascertained when the performance of the subjects were -1.5 standard deviations (SD) or below that of the age-, gender-, and education-adjusted norms in any of the CERAD-K-N neuropsychological tests.

This study was approved under the recommendations of the Institutional Review Board (IRB) of the Seoul National University Bundang Hospital (SNUBH, IRB approval number, B-2102-664-305), South Korea. All protocols and manuals were also approved by the IRB of the SNUBH.

A total of 53 vaccinees who received the BNT162b2 mRNA vaccine and 2 vaccinees who received the Oxford-AstraZeneca COVID-19 vaccine were recruited, and their peripheral blood sampling procedures were approved by the IRB of Seoul National University Hospital (SNUH, IRB approval number, 2102-032-1193). The details of the vaccination cohort data are described in a previous publication^15^.

### Blood sampling and preparation of total RNA from the peripheral blood mononuclear cells

At least 15 ml of peripheral blood was collected per patient in EDTA tubes (BD Vacutainer K2EDTA tubes, 367525)^35^. Peripheral blood mononuclear cells (PBMCs) and plasma were separated by density-gradient centrifugation using Ficoll (Cytiva Ficoll-Paque Plus, 17-1440-02) solution. After the PBMC suspension was resuspended in CELLBANKER (Amsbio CELLBANKER 2, 11914), the collected PBMCs were immediately cryopreserved and stored in a freezing container (Nalgene Mr. Frosty, C1562) for a week. The total RNA was isolated by TRIzol Reagent (Invitrogen TRIzol Reagents, 15596026) following the manufacturer’s instructions.

### Preparation of next-generation sequencing (NGS) library

For the NGS library preparation of BCR heavy chain sequences, genes encoding the variable region of the heavy chain (V_H_) at the 5’ forward region and the first constant region of the heavy chain (CH1) domain at the 3’ reverse region were amplified by specific PCR primers as reported previously^18^. All primer sequences are listed in Extended Table 2. In short, 1 *μ*g of total RNA was used as the input template for library preparation. Next, reverse transcription was performed using the SuperScript IV First-Strand Synthesis System (Invitrogen, 18091050) following the manufacturer’s instructions with CH1 gene-specific primers for five Ig heavy chain isotypes containing UMI (unique molecular identifiers) barcodes. The UMI barcodes consisted of 12 random nucleotides, 14 base pairs in total; NNNNTNNNNTNNNN, and partial Illumina adapters (No. 1 - 8). After synthesis of the first-strand cDNA, it was purified with SPRI beads (Beckman Coulter AMPure XP Reagents, A63882) at a 1:1.8 ratio, and the second-strand cDNA was synthesized using a KAPA Biosystems kit (Roche KAPA HiFi Hotstart PCR kit, KK2502) with V_H_gene-specific primers (No. 9 - 14; 95°C for 3 min, 98°C for 30 s, 60°C for 45 s, and 72°C for 6 min). The double-stranded cDNA was purified through SPRI beads at a 1:1 ratio, then amplified using the KAPA Biosystems kit and double primers containing Illumina adapters and index sequences (No. 15, 16; 95°C for 3 min, 25 cycles of 98°C for 30 s, 60°C for 30 s, 72°C for 1 min, and 72°C for 5 min). The PCR products were then subjected to electrophoresis on a 1.5% agarose gel for separation by DNA lengths and purified using the QIAquick gel extraction kit (QIAGEN, 28704) following the manufacturer’s protocols. The final NGS libraries were obtained using SPRI beads at a 1:1 ratio and quantified with a 4200 TapeStation System (Agilent Technologies, G2991BA). Libraries with a single peak of appropriate length were subjected to NGS analysis using the Illumina Novaseq6000 platform.

### NGS data processing

The raw NGS forward (R1) and reverse (R2) reads were merged, filtered based on Phred scores, and subclustered using UMI with paired-end read merger (PEAR) v0.9.10 and Clustal Omega v1.2.4 with default settings^36,37^. The sequence annotation process was completed using the IMGT (International Immunogenetics Information System) database^38^ to obtain the isotypes, and IgBLAST v1.17.1^39^ to obtain VJ genes, CDR1/2/3 sequences, and the number of mutations from the corresponding V genes, as previously described^18^. To exclude the possibility of contamination due to index swapping within Illumina flow cells between samples, reads with identical sequences including the UMI from different samples were removed on the basis that such an event is highly unlikely without contamination^40^. The combined NGS throughput of each sample is listed in Extended table 3. Sequencing reads that have survived through the entire process were considered to be productive BCR sequences. Files with less than 35,000 functional BCR sequences were excluded from the study.

### Selection of stereotypic AD patient-specific non-naïve BCR clonotypes

A stereotypic BCR clonotype is defined in this paper as a group of sequences sharing identical IGHV and IGHJ genes and the same CDR3 amino acid sequence while being present in two or more AD patients or vaccinees (Step 2 & 3, Fig 1c)^18^. Clonotypes which consist only of naïve sequences, those with the IgM or IgD isotype with one or no somatic mutation were excluded from further analysis (Step 1, Fig. 1c). Due to the potential risk of molecular contamination during the NGS library preparation stage, an additional strict *in silico* filtering step was performed. To exclude the possibility of molecular contamination during NGS library preparation or NGS, we confirmed that two or more AD patients or CG subjects with different BCR sequences could be identified for each clonotype (Step 4, Fig 1c).

### Sharing degree analysis of AD patient pairs

The similarity scores based on shared stereotypic AD-patient specific non-naïve BCR clonotypes at each TP were calculated as the cosine similarity of two vectors each representing a subject’s repertoire. The vectorization process involved utilizing one-hot encoding to create a vector with the length equal to the total number of cohort-specific and shared BCR clonotypes. The presence or absence of each clonotypes for the patient was encoded as either the value 1 or 0. The cosine similarity calculated as the inner product divided by the product of the size of each vector can be interpreted as the number of shared cohort-specific shared clonotypes in a patient pair divided by the square root of the product of the total number of cohort-specific shared clonotypes for each patient. The same analysis was performed on the CG.

### Construction of scFv phage display library and bio-panning

For the V_H_ gene, the V3-7/J1 VRLAEYFQN clonotype was chosen for further testing of its affinity against the Aβ42 peptide. Four of these BCR sequences were synthesized (Integrated DNA Technologies). All primer sequences are listed in Extended Table 2. For the construction of V_κ_/V_λ_ shuffled libraries, the total RNA from patients who possessed the V3-7/J1 VRLAEYFQN clonotype at TP2 (AD9, AD14 and AD34) were used to synthesize cDNA using the SuperScript IV First-Strand Synthesis System (Invitrogen) with gene-specific primers annealing to the C_κ/λ_ genes (No. 17 - 20). The cDNA was purified with SPRI beads (Beckman Coulter AMPure XP Reagents) at a 1:1.8 ratio, and the purified product was subjected to the first round of PCR synthesis using V_κ_/V_λ_ and J_κ_/J_λ_ gene-specific primers (No. 21 - 57; 95°C for 3 min, 4 cycles of 98°C for 1 min, 60°C for 1 min, 72°C for 1 min, and 72°C for 10 min). The PCR product underwent another purification step with SPRI beads at a 1:1 ratio, and the synthesized V_H_ genes and V_κ_/V_λ_ genes were amplified using a KAPA Biosystems kit (Roche) with the following primers (No. 58 - 61; 95°C for 3 min, 27 cycles of 98°C for 30 s, 65°C for 30 s, 72°C for 1 min, and 72°C for 10 min). Then, the amplified gene fragments of the V_H_ and V_κ_/V_λ_ were each subjected to electrophoresis on a 1.5% agarose gel and purified via DNA size using the QIAquick gel extraction kit (QIAGEN) following the manufacturer’s instructions. The amplified V_H_ and V_κ_/V_λ_ gene fragments (100 ng of each) from the 3 patients were mixed at an equal ratio to make a single pool of V_H_ and V_κ_/V_λ_ genes. The equally mixed pool of V_H_ and V_κ_/V_λ_ gene fragments was subjected to overlap extension PCR to generate scFv genes using a KAPA Biosystems kit (Roche) with overlap extension primers (No. 62, 63; 95°C for 3 min, 25 cycles of 98°C for 30 s, 65°C for 30 s, 72°C for 1 min 30 s, and 72°C for 10 min). The amplified scFv genes were then purified and cloned into a phagemid vector as described previously^41^. After electroporation into the bacterial cells and phage rescue, a phage display of human scFv library with 1.1 × 10^9^ colony-forming units was prepared^41^. The scFv library was subjected to four rounds of bio-panning against the human Aβ42 peptide (Tocris Bioscience Amyloid β-Peptide 1-42 human, 1428), as described previously^42,43,44^. Briefly, 2 *μ*g of the Aβ42 peptide was coated to the surface of the immunotube with a high binding surface (SPL Life Sciences, 43015) for 16 hours at 4°C, and blocked with 5 ml of 3% bovine serum albumin (BSA) in PBS (w/v). Then the scFv phage library (about 10^13^ phages) was incubated for 2 hours at room temperature. During the first round of panning, the tube was washed once with 5 ml of 0.05% (v/v) Tween 20 (Sigma-Aldrich, P1379) in phosphate-buffered saline (PBST). After each round of panning, the phages bound to the coated antigen were eluted and rescued for the next round of bio-panning. For the remaining rounds of panning, the number of washes was increased to three times. To select antigen-reactive scFv phage clones, phage clones from the titration plate of the last round of panning were randomly selected and subjected to phage ELISA, using the antigen-coated microtiter plates. Reactive scFv clones were analyzed through Sanger nucleotide sequencing using their phagemid DNA extracted from the individual phage clones, as described previously^18^. A recombinant scFv fusion protein tagged with human IgG1 Fc fragment (hFc) was expressed in the mammalian expression system and purified following the methods described previously^18^.

### Enzyme-linked immunosorbent assay (ELISA)

The reactivity of scFv-displayed phage clones and recombinant scFv-hFc fusion proteins were assessed by ELISA as described previously^18^. In short, the human Aβ42 peptide (Tocris Bioscience) was dissolved at a concentration of 100 nM in 50 *μ*𝓁 of 0.1 M sodium bicarbonate coating buffer (pH 8.6), then added to each well of 96-well microtiter plates (Corning, 3690) and incubated at 4°C for 16 hours. After the plates were blocked with 3% BSA in PBS (w/v) for 1 hour at 37°C, phage culture supernatant (diluted twofold) or recombinant scFv-hFc fusion proteins (serially diluted fourfold, 7 dilutions, starting from 250 nM to 61 pM) in blocking buffer were added to the wells of the microtiter plates, then incubated for at 37°C for 2 hours. Then, the plates were washed three times with 0.05% PBST (v/v). Either the horseradish peroxidase (HRP)-conjugated anti-M13 antibodies (Sino Biological, 11973-MM05T-H, 1:4,000) or HRP-conjugated rabbit anti-human IgG Fc antibodies (Invitrogen, 31423, 1:5,000) diluted in blocking buffer were added to the wells incubated with phages or recombinant proteins, respectively, and incubated at 37°C for 1 hour. After three washes with 0.05% PBST, 2,2′-azino-bis-3-ethylbenzothiazoline-6-sulfonic acid solution (Invitrogen, 002024) was added to the wells as the coloring substate for HRP. After the coloring reaction, absorbance was measured at 415 nm using a microplate spectrophotometer (Thermo Fisher Scientific Inc., Multiskan GO). All ELISAs were performed in duplicate.

### Statistical analysis

The P values were calculated using the student t-test unless otherwise specified during comparison of BCR repertoire characteristics. P-value < 0.05 was regarded as significant.

## Data availability

The raw sequencing data and processed data have been submitted to the NCBI GEO database under the accession number GSE242738.

## Acknowledgements

This research was supported by 1) the Korea Dementia Research Project grant through the Korea Dementia Research Center (KDRC), funded by the Ministry of Health & Welfare and Ministry of Science and ICT, Republic of Korea (grant number: HU20C0339), 2) the BK21 FOUR program of the Education and Research Program for Future ICT Pioneers, Seoul National University in 2022, 3) the Ministry of Science and ICT(MSIT) of the Republic of Korea and the National Research Foundation of Korea (NRF-2020R1A3B3079653) and 4) Hyundai Car Chung Mong-Koo Foundation.

## Additional information

Supplementary Information is available for this paper.

Correspondence and requests for materials should be addressed to J. Chung.

---

**Extended figure 1.**
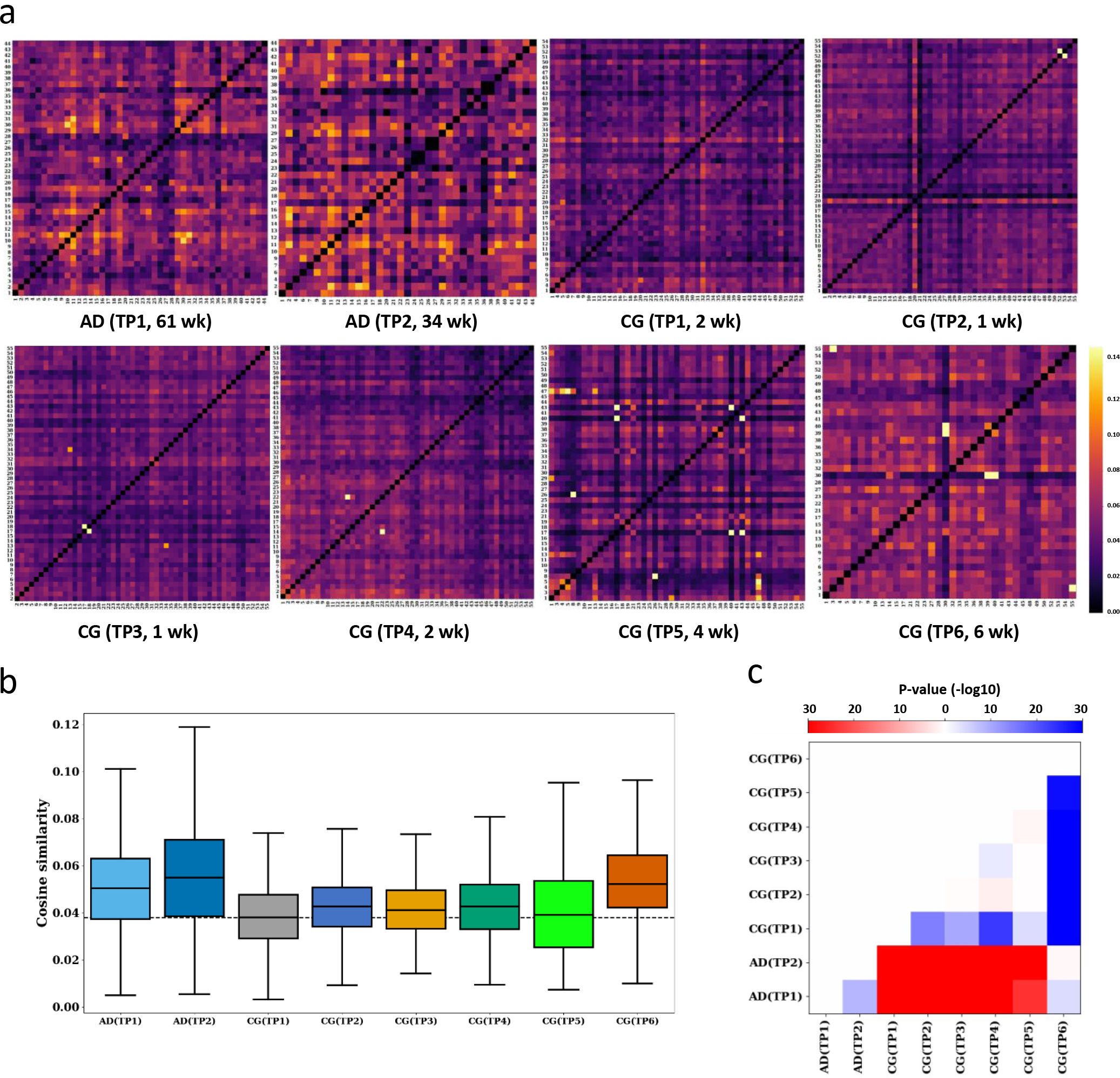
The sharing degree of stereotypic non-naïve BCR clonotypes among AD patient and CG subject pairs drawn, in which the clonotypes present in both groups were not filtered out. a, Heat maps show the sharing degree of stereotypic AD patient- or CG subject-specific non-naïve BCR clonotypes in AD patient or CG subject pairs. The cosine similarity between two one-hot encoded linear vectors representing the presence (1) or absence (0) of each stereotypic BCR clonotype. PB sampling TPs and their durations are denoted inside parentheses. b, Boxplots representing the distribution of cosine similarity for all AD patient or CG subject pairs at each PB sampling TP. The dashed line represents the median cosine similarity value of CG subject pairs at TP1. c, A heat map representing the P-values about the difference in their cosine similarity value in AD patient or CG subject pairs at every PB sampling TPs. The color of the heat maps indicate whether the median value on the y axis is larger (red) or smaller (blue) than that on the x-axis. The intensity of said colors is encoded by the P-value.

**Extended figure 2.**
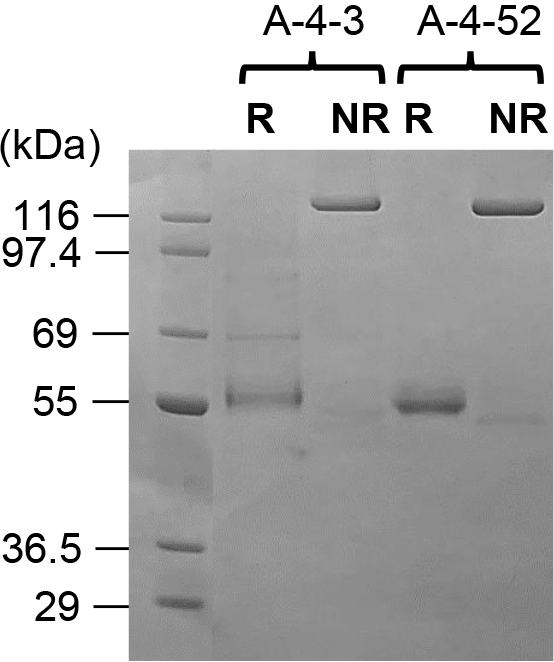
Expression of recombinant scFv-hFc fusion proteins. Aβ42-binding scFv clones were successfully expressed as recombinant scFv-hFc (human IgG1 Fc) fusion protein using a mammalian expression system. The expected protein size is 56 kDa in the non-reducing state (NR) and the dimeric protein is cleaved in reducing conditions (R) as single disulfide bond in the hinge region is broken.

